# Loss of CHMP2A implicates an ordered assembly of ESCRT-III proteins during cytokinetic abscission

**DOI:** 10.1101/2025.06.24.661003

**Authors:** Nikita Kamenetsky, Dikla Nachmias, Suman Khan, Ori Avinoam, Itay Hazan, Alexander Upcher, Natalie Elia

## Abstract

The ESCRT machinery mediates membrane remodeling in fundamental cellular processes including cytokinesis, endosomal sorting, nuclear envelope reformation, and membrane repair. Membrane constriction and scission is driven by the filament-forming ESCRT-III complex and the AAA-ATPase VPS4. While ESCRT-III-driven membrane scission is generally established, the mechanisms governing the assembly and coordination of its twelve mammalian isoforms in cells remain poorly understood. Here, we examined the spatial organization and interdependence of ESCRT-III subunits during mammalian cytokinetic abscission by depleting CHMP2A, a core ESCRT-III component. Using live cell imaging, structured illumination microscopy (SIM) and correlative light-electron microscopy (CLEM), we show that CHMP2A knockout cells display a significant delay—but not failure—in abscission, accompanied by distinct mislocalization phenotypes across ESCRT-III subunits. While IST1 and CHMP2B were minimally disrupted, CHMP4B, CHMP3, and CHMP1B display progressively severe organization defects at the abscission site. Dual- protein imaging reveals disrupted coordination between ESCRT-III subunits in individual CHMP2A-deficient cells, supporting an ordered assembly of ESCRT-III subunits in cytokinetic abscission. Together, our findings provide the first in vivo evidence for hierarchical assembly of ESCRT-III subunits during ESCRT-mediated membrane remodeling and identify CHMP2A as a key organizer of ESCRT-III architecture essential for timely membrane abscission.

## INTRODUCTION

The endosomal sorting complex required for transport (ESCRT) is an evolutionarily conserved machinery that facilitates the remodeling and scission of membranes. ESCRT are required for various processes in cells, including multivesicular body (MVB) biogenesis, viral budding, membrane repair, nuclear envelope sealing, nuclear pore complex quality control, and abscission of the intercellular membrane tube during cytokinesis [1–3]. Despite progress in identifying proteins involved in membrane shaping and remodeling, the exact mechanisms by which ESCRTs bend and cut membranes have yet to be resolved.

The ESCRT machinery is composed of five different subfamilies: ESCRT 0, I, II, III, and the AAA ATPase VPS4. ESCRT-III and VPS4 constitute the minimal ESCRT module required for membrane fission. According to the current model, recruitment of early ESCRTs (ESCRT 0- II) to the membrane, facilitates the recruitment of late ESCRT-IIIs. ESCRT-IIIs then self- polymerize into helical filaments on the inner side of the membrane. These filaments, in the presence of VPS4, facilitates membrane constriction and fission [3].

Twelve mammalian ESCRT-III subunits have been identified in mammalian cells: CHMP1A, CHMP1B, CHMP2A, CHMP2B, CHMP3, CHMP4A, CHMP4B, CHMP4C, CHMP5, CHMP6, CHMP7, and IST1 (CHMP8). The rationale for the emergence of multiple ESCRT-III genes is still unclear. *In-vitro*, diberent ESCRT-IIIs assemble into distinct polymeric shapes including; tubes, spirals, cones, and coils, with varying diameters and thicknesses [3–6]. Moreover, the yeast homolog of CHMP4B, Snf7, polymerize on liposomes into a 2D single-strand spiral when present alone and addition of other ESCRT-III subunits was found to abect the thickness and dynamics of the Snf7 spiral [7]. Finally, in the presence of VPS4, the Snf7 polymer was able to exchange specific ESCRT-III subunits [7–9]. Together these in vitro findings suggested a model in which ESCRT-IIIs, promote membrane constriction by organizing into dynamic polymers, that exchange their subunit composition, facilitating a transition from large to small diameter spiral. However, this model has yet to be demonstrated in cells.

Cytokinetic abscission, the last step of cell division, in which the intercellular bridge that connects the two daughter cells is resolved, represent an ideal system to study the spatiotemporal recruitment of ESCRTs in physiological context [10] (Fig. 1A). Constriction and fission of a ∼1 μm-thick intercellular bridge, was shown to be driven by the ESCRT machinery in a well-orchestrated and regulated process [11–14]. Microscopic analysis of cytokinetic abscission, using Structured Illumination Microscopy (SIM) and high-sensitivity temporal imaging of fluorescently labeled ESCRT proteins (obtained by confocal spinning disk microscopy), revealed that ESCRTs I-III and VPS4 are recruited sequentially to the rims of the dark zone (DZ), located at the center of the intracellular bridge, forming ring-like structures [11, 15]. Proteins of the ESCRT-III complex, were shown to initially assemble into ring-like structures, and then polymerize to elongated helical structures with reduced diameters, previously documented by STORM [16]. While super resolution studies mainly focused on CHMP4B and IST1, other ESCRT-IIIs were found to localize to the intercellular bridge, suggesting their potential involvement in bridge constriction [12, 17]. Yet, the relative spatio-temporal recruitment of these proteins and their contribution to abscission has yet to be resolved.

**Figure 1.**
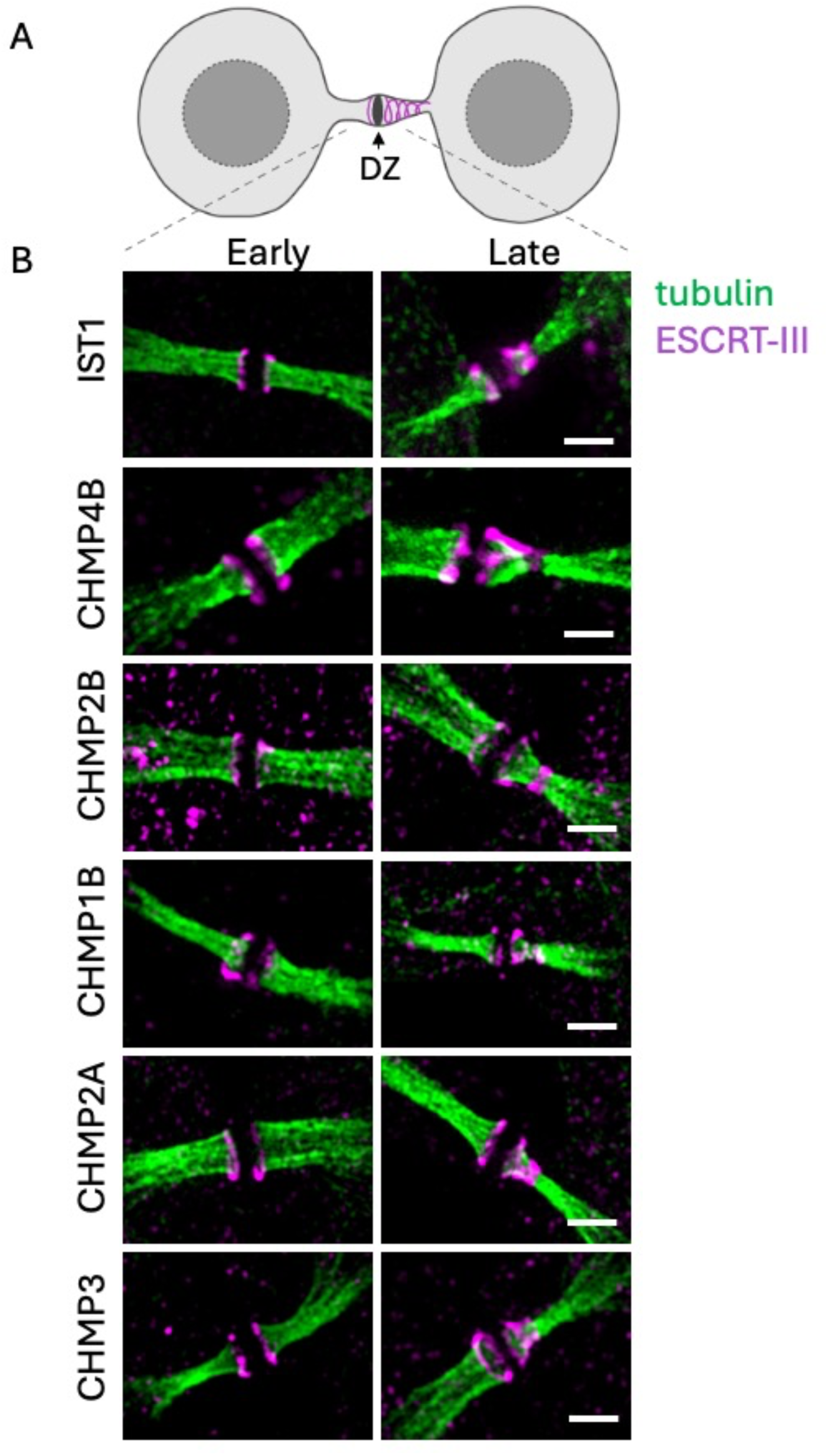
ESCRT-III Proteins Exhibit a Conserved Assembly Pattern During Membrane Constriction and Abscission. (A) Schematic illustration of cells undergoing abscission. Cells are connected via an intercellular bridge. A dense area called the dark zone (DZ) is located at the center of the bridge (black). ESCRT-IIIs organize into high-ordered structures on both sides of the intercellular bridge (magenta). Left of the DZ, early ring-like organization; right of the DZ, late spiral-like organization. (B) Representative maximum projection SIM images of synchronized HeLa cells, stained with antibodies against α-tubulin (green), and the indicated ESCRT-IIIs (magenta). Shown are intercellular bridges at early (left) and late (right) stages. In early intercellular bridges, all ESCRT-IIIs exhibited the typical organization of two rings located at the rims of the DZ. In late intercellular bridges, all ESCRT-IIIs exhibited asymmetric structural organization with one side (left of DZ) exhibiting ring-like structure and the other side (right of DZ) exhibiting elongation toward the center of the bridge that is accompanied with bridge constriction. Data was reproduced in at least 3 independent experiments. Scale = 1 μm.

The ESCRT-III protein CHMP2A, was shown to be a crucial subunit of the ESCRT-III machinery both *in-vitro* and in cells. *In-vitro*, CHMP2 was found to mediate the transition from large to small diameter spiral transition, during VPS4-mediated subunits exchange [7]. Based on the biochemical and structural studies, CHMP2A was found to pair with CHMP3, to form capping assemblies, that interact with membranes through electrostatic forces, which potentially drive final fission of the membrane [18–20]. In cells, CHMP2A was found to be involved is most ESCRT mediated cellular functions, including; cytokinetic abscission, membrane wound repair, nuclear envelop sealing and autophagy [2]. Moreover, viruses, such as HIV-1, exploit CHMP2A for viral budding, enhancing their infectivity [18, 21]. Dysregulation of CHMP2A, was also implicated in a range of pathologies. Mutations or altered expression of CHMP2A contributed to neurodegenerative disorders, like Alzheimer’s and Parkinson’s diseases, potentially by impairing the degradation of toxic protein aggregates, through disruption of phagophore closure [22, 23]. Tumor cell resistance to immune system attacks, was also found be regulated by CHMP2A [24]. These mechanistic and pathological roles of CHMP2A points to its fundamental importance in ESCRT-mediated membrane remodeling.

In this study, we set out to determine the interplay between ESCRT-IIIs during abscission. Given the fundamental role proposed for CHMP2A, we generated CHMP2A KO cells and tested whether absence of CHMP2A abects the organization of diberent ESCRT-IIIs at the intercellular bridge, and the abscission process. Using SIM of endogenous ESCRT components and correlative Light-EM microscopy (CLEM), we found that in WT cells all ESCRT-IIIs exhibit an overlapping organization pattern at the bridge. Diberential organization was observed for ESCRT-IIIs in CHMP2A KO cells, with some ESCRT-IIIs exhibiting a highly severe organization phenotype, while others were only mildly abected. Notably, CHMP2A KO cells could still undergo abscission, albeit at a slower rate, inferring to the robustness of the ESCRT system. Our findings provide the first in vivo evidence to support a hierarchical assembly of ESCRT-IIIs during abscission, contributing to a mechanistic understanding of ESCRT-driven membrane abscission.

## RESULTS

Several studies described the arrival of various ESCRT-IIIs at the intercellular bridge [11, 25, 26]. Yet, most studies, relied on visualization of overexpressed fluorescent ESCRT-III variants, which may differ from endogenous protein behavior. To systematically investigate the organization of different ESCRT-IIIs, at the intercellular bridge, at native conditions, we employed antibody staining followed by SIM imaging. We found that IST1, CHMP4B, CHMP2B, CHMP1B, CHMP2A, and CHMP3, all follow the typical organization described for ESCRT-IIIs at the intercellular bridge, i.e. an initial large diameter ring at early bridges, and an elongated cone structure at late bridges [12, 16] (Fig. 1B). These results are consistent with previous studies, showing in high resolution that under endogenous conditions, most ESCRT-IIIs arrive at the intercellular bridge, and organizes into high-ordered structures [16, 17].

Previous *in-vitro* studies inferred a critical role for CHMP2A in the transition of the ESCRT-III spiral to a constricted state. To test this notion in a cellular setup, we generated a CHMP2A knock-out (KO) HeLa cell line, using CRISPR genome editing system. We verified CHMP2A loss by western blot (Fig. 2A). Of note, expression of other ESCRT-IIIs was not affected in the KO cells (Fig. S1). Initial characterization of CHMP2A KO cells, revealed a ∼two-fold increase in overall cell size, which was proportional to nuclear size, maintaining the cytoplasm-to-nucleus ratio obtained in WT cells (Fig. S2). Consistently, the diameter of the intercellular bridge was larger in CHMP2A KO cells, compared to WT cells (2.25 ± 0.17 μm vs 1.85 ± 0.17 μm for WT and KO cells, respectively) (Fig. 2B). Live cell imaging of dividing CHMP2A KO cells, showed that cytokinetic abscission is considerably delayed. CHMP2A KO cells completed abscission at an averaged time of ∼154 minutes, compared to ∼80 minutes in WT cells (Fig. 2C-D). In contrast to other ESCRT depletion conditions that resulted delayed abscission [16], the observed abscission delay in CHMP2A KO cells was not accompanied with abscission failure and all cells imaged successfully completed abscission. The membrane constriction profiles of intercellular bridges undergoing abscission were similar in WT and CHMP2A KO cells, suggesting that, in the absence of CHMP2A, abscission progresses normally albeit at a slower rate (Fig. 2D).

**Figure 2.**
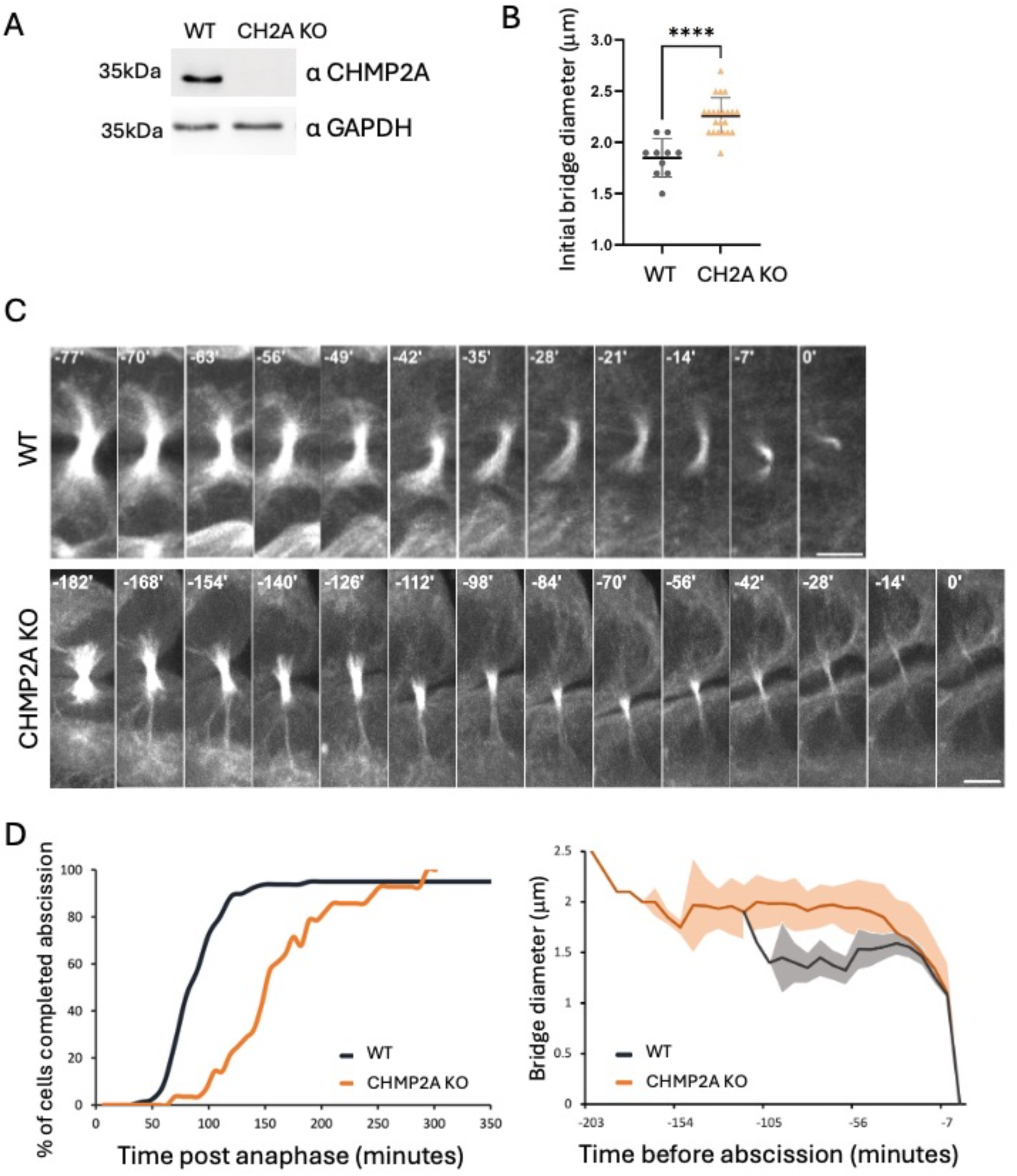
Loss of CHMP2A delays abscission but does not lead to abscission failure. (A) Western blot analysis of CHMP2A expression levels (32 kDa) in WT and CHMP2A KO cells. Anti-GAPDH (bottom panel) was used as loading control. Equal protein amounts were loaded. (B) Initial intercellular bridge diameter, as determined based on GFP-tubulin intensity at the end of anaphase in HeLa WT (black, n=10) and CHMP2A KO (orange, n=21) using a line intensity profile. Statistical significance was assessed using a two-tailed Mann–Whitney U test. *p < 0.001 (\*\*\*\**). (C) Representative maximum projection confocal time-lapse images, of HeLa cells transfected with GFP-tubulin (grey) undergoing abscission. WT, top panel; CHMP2A KO, bottom panel. Cells were imaged at 7-minute intervals. Time = 0 was set as the time of bridge scission, based on GFP-tubulin. Scale = 5 µm. (D, left) Cumulative graph depicting the percentage of cells that completing abscission over time in WT (black, n=76) and CHMP2A KO (orange, n=28) cells. time = 0 was set anaphase at onset. (D, right) Intercellular bridge diameter measured for individual bridges over time based on GFP-tubulin intensity in cells undergoing abscission using a line intensity profile. WT, black, n=10; CHMP2A KO, orange, n=21 cells. Mean ± SD is plotted. Data was obtained from at least three independent experiments.

To investigate whether the delayed abscission observed in CHMP2A KO cells arises from altered assembly of ESCRT-IIIs at the intercellular bridge, we followed ESCRT-IIIs localization at late-stage intercellular bridges. First, we immunolabeled cells with CHMP4B antibodies and imaged late-stage intracellular bridges of dividing cells by SIM (Fig. 3A). While ∼40% of the cells exhibited the typical organization, described for ESCRT-III at the intercellular bridge i.e., a large-diameter cortical ring at the rims of the DZ, and a helical filament, stretching from the edge of the DZ to the abscission site (Normal) (see also Fig.1B), ∼60% of the cells exhibited defective organization (Fig. 3B-C). We classified the defective organization phenotypes, into three categories based on their severity. The least severe phenotype (Assembled) was manifested by proper localization at the DZ, with abnormal / defective ring-assemblies. The second phenotype (DZ Localization) was characterized by accumulation of CHMP4B at the DZ, without assembly into high-ordered structures. The most severe phenotype (No DZ localization) was marked by a complete failure of CHMP4B to localize to the DZ; instead, the protein appeared in a string-like pattern along the intercellular bridge (Fig. 3B). This latter phenotype could also readily be detected by confocal microscopy (Fig. S3). Therefore, loss of CHMP2A leads to defects in CHMP4B organization at the intercellular bridge.

**Figure 3.**
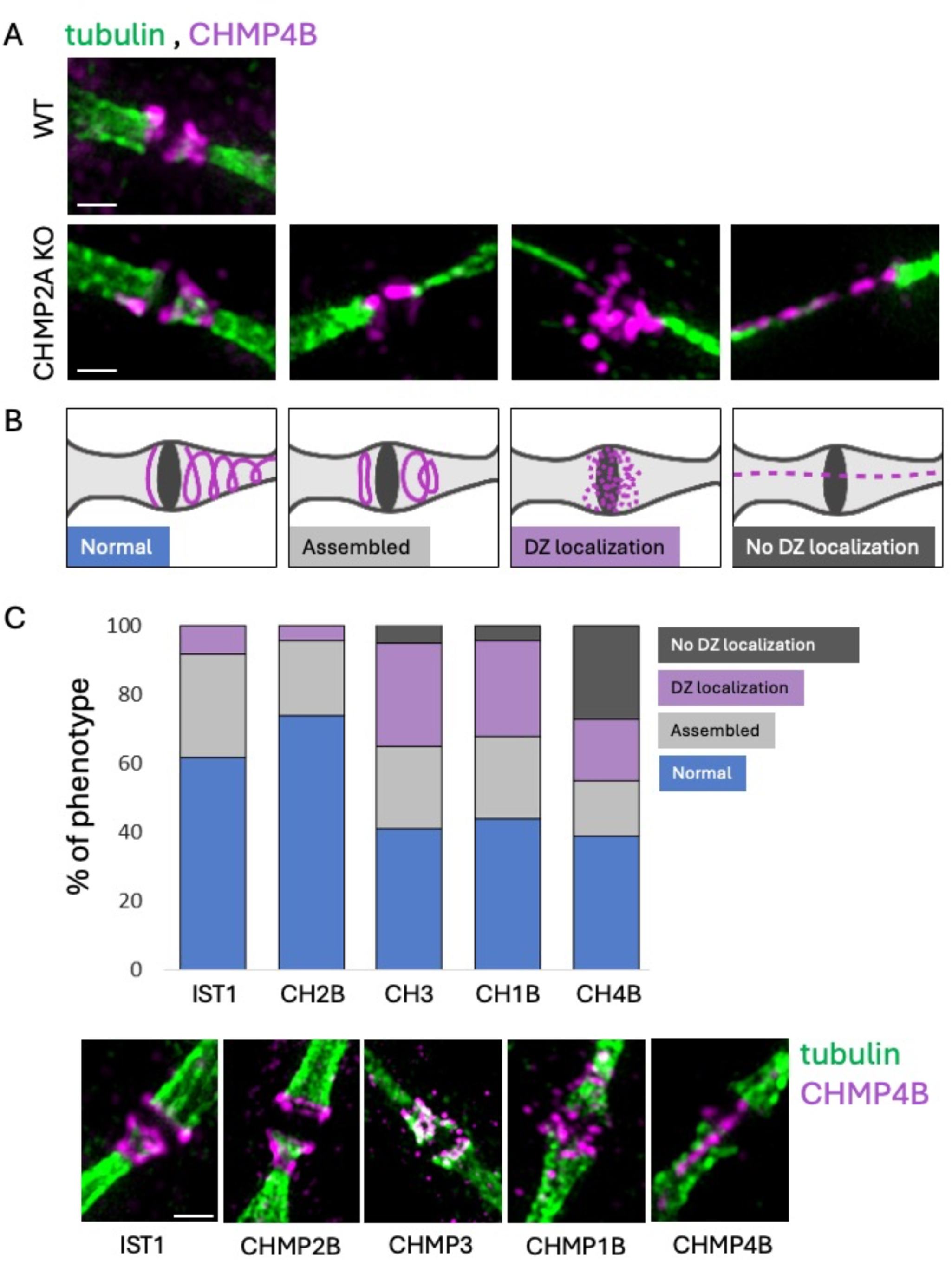
Chmp2a depletion alters the assembly and localization patterns of escrt-iii proteins at the intercellular bridge. (A) Representative maximum projection SIM images of synchronized HeLa cells. WT, top; CHMP2A KO, bottom) stained with α-tubulin (green) and α-CHMP4B antibodies (magenta). Various localization defective phenotypes observed at the intercellular bridge upon CHMP2A depletion are shown. (B) Schematic illustration of the observed phenotypes: Typical localization observed in WT cells (Normal); ESCRT-IIIs adopted a high ordered structure, but with abnormal morphology (Assembled); ESCRT-IIIs localized at the intracellular bridge but failed to organize in high ordered structures (DZ localization); ESCRT-IIIs did not organize at or near the DZ. Instead, the proteins adopted a string-like pattern along the intercellular bridge (No DZ localization). (C) Relative abundance of the observed phenotypes, as obtained for different ESCRT-IIIs in CHMP2A KO cells. Synchronized HeLa CHMP2A KO cells were stained and imaged (as in A), using specific antibodies for ESCRT-IIIs, as indicated. Top: quantification of the relative abundance of the observed phenotypes. Bottom: Representative maximal projection SIM images at late stages of abscission, demonstrating the various phenotypes. IST1, n=90; CHMP2B, n=25; CHMP1B, n=29; CHMP4B, n=39. Statistical significance was calculated using Fisher’s exact test by comparing the percentage of normal vs abnormal localization patterns for each protein in WT and KO cells. Calculated p values were < 0.0001 for all proteins analyzed. Data was obtained from at least two independent experiments. Scale = 1 μm.

Interestingly, expanding this analysis to additional ESCRT-IIIs, resulted in similar phenotypic defects, albeit at different ratios. In CHMP2A KO cells, CHMP3, CHMP1B, and CHMP4B exhibited abnormal organization in over 50% of cells, with CHMP4B exhibiting the most pronounced defects (27% of the cells exhibiting the ‘No DZ localization’ pattern) (Fig. 3C). This was not case for IST1 and CHMP2B, which displayed normal localization in the majority of cells (62% and 74%, respectively), and did not exhibit ‘No DZ localization’ phenotype in any of the CHMP2A KO cells we imaged. These results suggest that absence of CHMP2A, differentially affects the organization of different ESCRT-IIIs at the intracellular bridge. The severity of the phenotype appears to follow an order, in which CHMP4B is the most severely affected, followed by CHMP1B, CHMP3, and finally IST1and CHMP2B, which were least affected by CHMP2A depletion.

To examine, whether the phenotypes observed using immunostaining, can also be detected at the ultrastructural level, we developed a correlative light and electron microscopy (CLEM) protocol, and imaged intercellular bridges exhibiting specific phenotypes (Fig. 4). HeLa cells were prepared both for fluorescence microscopy (FM) of antibody tagged components as well as for electron microscopy (EM; see methods). Ultimately, we were able to map and image individual intercellular bridges by FM, and by EM, allowing correlation of specific protein identity and ultrastructural data (see Methods section). To validate the approach, we initially stained WT cells with anti-IST1 antibodies, which we have previously optimized for ESCRT-III staining at intercellular bridge, and imaged intercellular bridges with Transmission electron microscopy (TEM) imaging of 350 nm thick slices [12, 16]. IST1 fluorescence was successfully preserved at the intercellular bridge, showing the typical organization described for ESCRT-III at late intercellular bridges (Fig. 4A). The corresponding electron micrograph confirmed that the observed structure was a late-stage bridge, exhibiting the microtubules constriction at the so called ‘constriction site’ [12]. Additionally, a dense structure, which was previously associated with the filaments of ESCRT-III assemblies, was clearly seen on the same side of the bridge, where IST1 localization was detected, strongly validating our CLEM protocol (Fig. 4A). Notably, this data represents the first direct indication that the helical filaments observed by EM are associated with the late-bridge organization described for ESCRT-III localization at the intercellular bridge.

**Figure 4.**
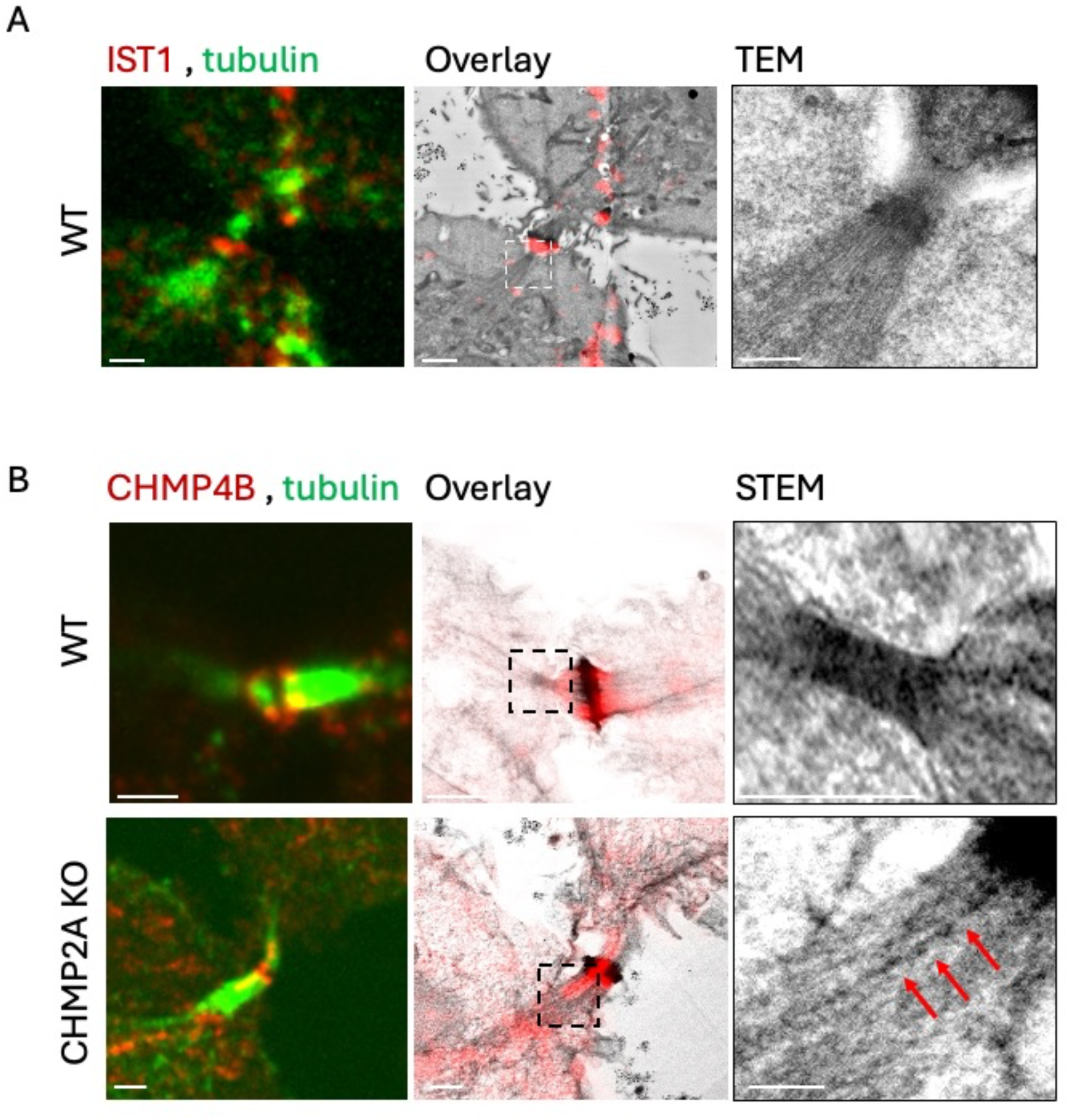
Correlative Light-EM supports the abnormal organizations detected for ESCRT-III at intercellular bridges of CHMP2A KO cells. (A-B) HeLa WT and CHMP2A KO cells were synchronized, stained and subjected to CLEM protocol as described in Methods section. (A) Cells were stained with α–tubulin (green) and α–IST1 antibodies (red), and imaged by spinning disk confocal followed by TEM on 350 nm-thick sections. (B) Cells were stained with α–tubulin (green) and α- CHMP4B antibodies (red) and imaged by spinning disk confocal followed by STEM on 1000 nm-thick sections. For each panel: Left, fluorescence image; middle, overlay of ESCRT-III protein and EM image; right, zoomed-in EM image. Red arrows in the CHMP2A KO STEM panel (B bottom, right) highlight string-like electron-dense structures that correlate with the CHMP4B signal observed by fluorescence microscopy. Shown are representative datasets. Scale left, middle panels = 2 µm, right panels = 1 µm.

Next, we examined the ultrastructural organization of intercellular bridges, displaying normal CHMP4B localization in WT cells. To capture the entire intercellular bridges, one-micron thick slices were imaged using Scanning Transmission Eelectron Microscopy (STEM) for subsequent tomography. We found that intercellular bridges, exhibiting the late-bridge localization pattern for CHMP4B by fluorescence, obtained the dense helical structure associated with ESCRT-III spiral on the same side of the bridge, by EM (Fig. 4B). This result further validates our correlative assay, and reinforce the spatial link between ESCRT-IIIs (by FM) and helical spirals (by EM) at the intercellular bridge (Fig. 4B). Lastly, we examined the ultrastructural phenotypes in CHMP2A KO cells, that exhibited the most severe ‘No DZ localization’ phenotype. Cells that exhibit the string-like localization of CHMP4B (by FM) did not obtain the high density at the periphery of the bridge seen in late intercellular bridges of WT cells (by EM). Instead, string-like densities were found along the bridge (arrows in Fig 4B), confirming this phenotype at the ultrastructural level. Collectively, this data suggests, that under normal conditions, extension of ESCRT-IIIs toward the periphery of the cell, is accompanied by the dense helical filamentous structure seen by EM. In cells that lack CHMP2A, CHMP4B often fails to integrate into the characteristic high-ordered ESCRT-III assemblies described in abscission, potentially leading to the delayed abscission observed in these cells.

The above results indicate that CHMP2A depletion induces differential localization phenotypes among ESCRT-IIIs. To further investigate this, we co-labeled pairs of ESCRT-IIIs in individual WT and CHMP2A KO cells and analyzed their localization patterns relative to one another by SIM, in late intercellular bridges. IST1, which exhibited a relatively normal organization in most CHMP2A KO cells (>60% Fig. 3C), was used as a reference for the analysis. In WT cells, all tested ESCRT-IIIs (IST1, CHMP1B, CHMP2B, CHMP3 and CHMP4B) co-localized, forming overlapping ESCRT-III ring-like / filamentous structures, supporting the co-organization of these ESCRT-IIIs. In contrast, CHMP2A KO cells, occasionally exhibited distinct localization phenotypes within the same cell (i.e, one ESCRT-III protein is properly localized while the other is impaired) (Fig. 5, Fig. S4). To quantify this effect, we specified the localization pattern for pairs of ESCRT-IIIs at individual intercellular bridges, using the four categories described above (’Normal’, ’Assembled’, ‘DZ localization’, ‘No DZ localization’) and plotted the data in a similarity matrix (Fig. 5B). This analysis revealed that in the same cells, ESCRT-IIIs are differentially affected by loss of CHMP2A, strongly suggesting an ordered assembly of ESCRT-IIIs at the intercellular bridge.

**Figure 5.**
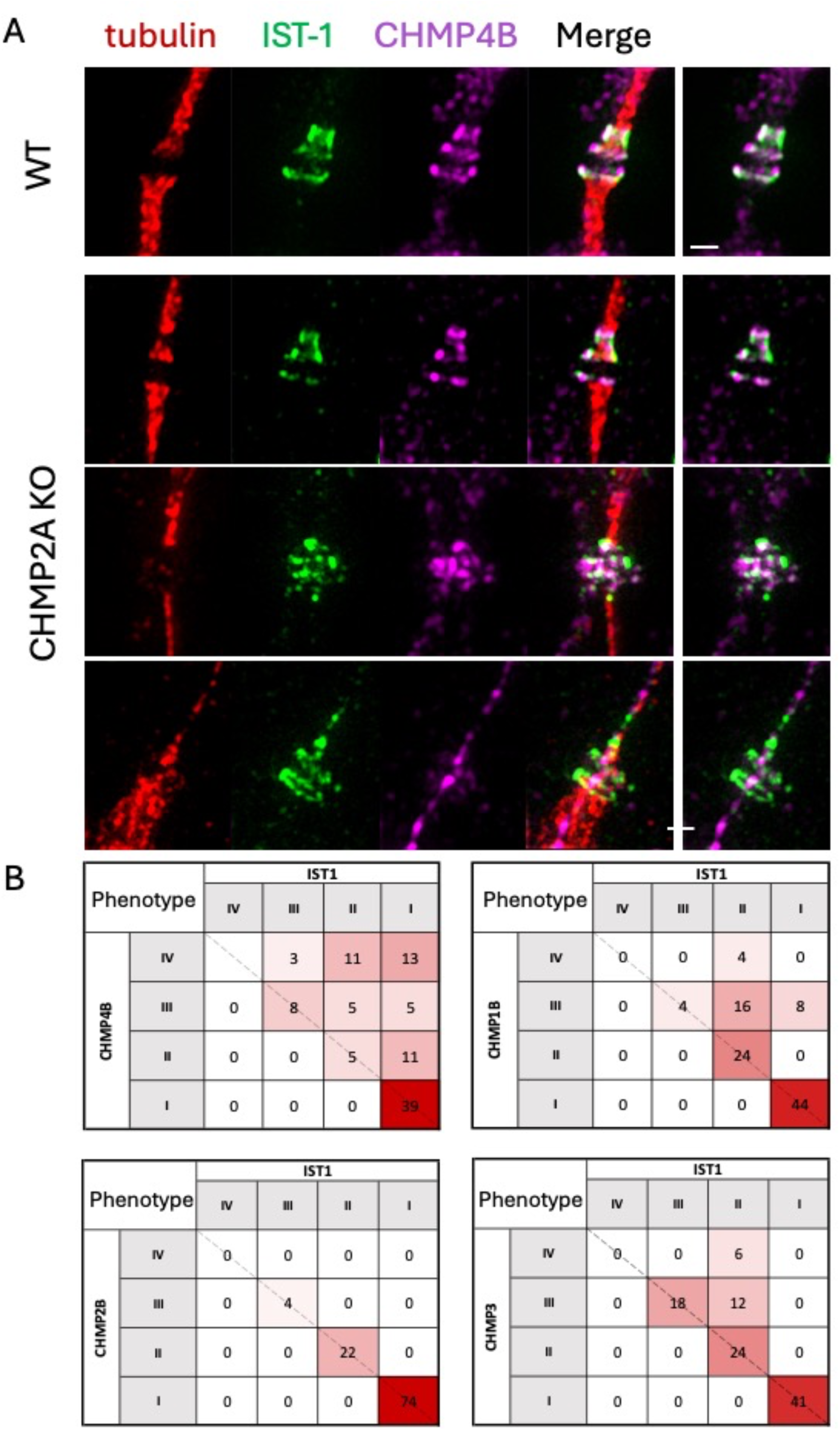
Differential localization patterns observed for ESCRT-III protein pairs in individual CHMP2A KO cells. Representative SIM images of synchronized HeLa cells (WT and CHMP2A KO) (A) HeLa WT (top panel) or CHMP2A KO cells (three bottom panels) were labeled with α– tubulin (red) and α-IST1 antibodies (green) and α-CHMP4B (magenta, third). Overlayed images are presented in fourth panels. Right panels, overlaid images of ESCRT-III proteins. Note the differential pattern observed for IST1 and CHMP4B in the bottom panel. Scale = 1 µm. (B) Percentages similarity matrix of the phenotypes described in Fig. 3 (I, Normal; II, assembled; III, DZ localization, IV, No DZ localization) as observed for IST1 with either CHMP4B, CHMP1B, CHMP3 or CHMP2B, as indicated. Red to white color-coding, indicates high to low frequencies, respectively. Values indicate Percentages. Representative images are shown in Fig. S4. n (number of cells) = IST1:CHMP4B (n=38), IST1:CHMP3 (n=17), IST1:CHMP1B (n=25), IST1:CHMP2B (n=27).

Among all ESCRT-III protein pairs tested, the highest proportion of cells showing similar localization patterns for two ESCRT-IIIs, was observed when both proteins were properly assembled at the intercellular bridge (‘Normal’/’Normal’ in all pairs). Overlapping defective localization patterns, were also observed in all pairs, as indicated by the percentages of cells along the diagonal in each similarity matrix. Abnormal IST1 localization, was always associated with abnormal localization of the second ESCRT-III protein tested, suggesting that IST1 is upstream to all the other ESCRT-IIIs we examined. Notably, the percentage of pairs with different localization phenotypes, between IST-1 to 2^nd^ ESCRT-III tested, were different between pairs. For example, IST1:CHMP4B pairs, exhibited the highest number of differential phenotypes (48%), while no instances of differential phenotypes were detected for IST1:CHMP2B pairs (100% of the cells imaged were along the diagonal of the matrix; Fig. 5B and Fig. S5 A-C). These findings suggest that the loss of CHMP2A disrupts the coordinated assembly of certain ESCRT-III components, while others remain unaffected. Moreover, the hierarchy of localization phenotypes, observed for different ESCRT-IIIs may infer to the order of ESCRT-III assembly at the intercellular bridge — where CHMP4B is the most affected, followed by CHMP1B, CHMP3, CHMP2B, and IST1. Proteins most affected by the absence of CHMP2A, such as CHMP4B, are likely to assemble downstream to CHMP2A while components such as CHMP2B and IST1, which are least affected, may function upstream of CHMP2A arrival and play a role earlier in the ESCRT-III assembly process. This dependency hierarchy underscores the central role of CHMP2A in orchestrating the spatial and temporal regulation of ESCRT-III assembly during cytokinetic abscission.

## DISCUSSION

In this study, we provide a systematic analysis of the localization pattern, of six different ESCRT-IIIs at the intercellular bridge of dividing mammalian cells. We found that depleting one of the ESCRT-IIIs, CHMP2A, differentially affect the organization of ESCRT-IIIs at the bridge. Loss of CHMP2A can affect the organization of two ESCRT-IIIs distinctively at the same bridge. This differential disruption in ESCRT-III localization was accompanied by delayed abscission, suggesting, that the native ESCRT-III structure formed at the intercellular bridge, is complex and rely on coordinated assembly of multiple ESCRT-IIIs, to efficiently execute its function. Nevertheless, the system is robust enough to ultimately complete the process when this coordination is disrupted.

Our phenotypic mapping of CHMP2A KO cells, showed that different ESCRT-IIIs are differentially affected. IST1 and CHMP2B exhibited mild defects while CHMP4B exhibited the most severe localization failure. Our results support a model, in which ESCRT-IIIs are orderly organized at the intercellular bridge and that CHMP2A, is required for this organization. Notably, the overall expression level of all ESCRT-IIIs tested was not significantly affected by CHMP2A KO suggesting that the localization phenotypes are not an outcome of a compensatory mechanism, that is induced by loss of CHMP2A (Fig. S1). Following this scenario, IST1 and CHMP2B are forming complexes, which are mostly independent of CHMP2A, while organization at the bridge of CHMP3, CHMP1B and CHMP4B rely on CHMP2A. This can suggest a sequential organization of ESCRT-IIIs at the intercellular bridge or the formation of several ESCRT-III high-order complexes, each comprising different subunits.

The findings that IST1 is only minimally affected by loss of CHMP2A, are aligned with our previous data, showing that IST1 is able to organize, in ring-like structures, at the intercellular bridge, upon siRNA depletion of the ESCRT-IIIs; CHMP4B, CHMP2B and CHMP1A [16]. Therefore, the ability of IST1 to organize into high-ordered structures at the intercellular bridge, appears to be less sensitive to the presence of other ESCRT-IIIs at the bridge. Notably, IST1 was originally identified as a crucial component of cytokinetic abscission [27, 28]. Collectively, these findings suggest that IST1 arrives and assembles early at the bridge, and independently of other essential ESCRT-IIIs such as CHMP2A and CHMP4B. CHMP4B, on the other hand, was shown to be most affected by CHMP2A loss. Interestingly, in more than 25% of the bridges, CHMP4B was found to reside in string-like structures located along the intercellular bridge, seen both by fluorescence and by EM. These structures may correspond to the recently reported vesicles that transports CHMP4B along microtubules in the intercellular bridge [29], suggesting that in the absence of CHMP2A, CHMP4B is delivered to the intercellular bridge, but accumulates in vesicle carriers instead of integrating into the ESCRT-III filament.

Previous EM-tomography and cryo-X-ray tomography data reported the presence of three parallel ESCRT-III fibers in the helical filament that reside at the intercellular bridge [12, 30]. The differential organization of ESCRT-III at the bridge described here may support a model in which each fiber is comprised of different ESCRT-III components and hence is differentially affected by loss of CHMP2A. Future studies, using higher spatial resolution in light microscopy, and employing our dual ESCRT-III labeling assay, may allow resolution of different ESCRT-III proteins at the bridge. Interestingly, in the archaeal ESCRT-III homologues CDV system, a set of three temporally separated rings comprising of different ESCRT-IIIs were found to drive cell division, suggesting a well-orchestrated ordered assembly of ESCRT-III filaments in prokaryotes [31, 32]. *In-vitro*, an ordered assembly of ESCRT-IIIs was described, in the presence of VPS4 [7]. Consistent with our findings, CHMP2A was found to mediate the transition between ESCRT-III monomers, in vitro, by interacting with both the early and late assembled ESCRT-IIIs [7]. Notably, while in the *in-vitro* setup, IST1 was the last to assemble forming the most constricted spiral, our data *in- cell*ule, suggests that IST1, is upstream to most other ESCRT-IIIs including CHMP2A, at least in the context of abscission. This difference could stem from differences between the *in-vitro* and *in-vivo* systems or may elute to the dynamic nature of the ESCRT system, which can assemble into different structures, holding different ESCRT-III compositions, and perhaps biophysical properties under different contexts or membrane topologies [3]. Nevertheless, the demonstration of ordered assembly of ESCRT-III proteins in vitro and in both prokaryotes and eukaryotes suggest that this is an evolutionary conserved property for ESCRT-III proteins, which is needed for driving ESCRT function.

## METHODS

### Cell culture and transfection

HeLa cells were grown in DMEM supplemented with 10% fetal bovine serum (FBS), 2mM glutamine, 10,000 U/mL penicillin, 10 mg/mL streptomycin at 37°C and 5% CO2.

Transfection was carried out by using Lipofectamine 2000 (Life Technologies), PolyJet (SignaGen Laboratories) or JetPrime transfection kit (Polyplus), according to manufacturer’s guidelines.

### Genome editing

For CRISPR/Cas9-mediated gene disruption, guide RNAs (gRNAs) targeting CHMP2A were sub-cloned into the lentiCRISPR plasmid (Addgene, #49535). Following transfection and puromycin selection, single clones were isolated and expanded. To confirm the efficacy of protein knockouts, western blot analysis was performed. The gRNA sequences employed were as follows:

5’- TGCGCAAGTTTGTATTGATG-3’

5’- CACTCGACAGCTCATCTGTT-3’

### Cell Synchronization

Double thymidine block was performed as previously described [ref inna cell reports]. In short, HeLa cells were plated at 10% confluency on an Ibidi Glass Bottom Dish 35 mm (Martinsried, Germany) and treated with 2 mM thymidine (Sigma, T1895-1G) for 18 hours to induce the first block. After incubation, cells were washed once with PBS and grown in fresh media for 9 hours. Cells were then supplemented for an additional 15 hours with 2 mM thymidine, washed once with PBS, and grown in fresh media for 10.5-12.5 hours to enrich the percentages of intercellular bridges.

### Western Blot

HeLa cells were lysed in RIPA lysis buffer [150 mM NaCl, 1% NP-40, 0.5% deoxycholate, 0.1% SDS, 50 mM Tris (pH 8.0)], supplemented with complete protease inhibitor (Roche Diagnostics) for 30 min at 4 °C. Total protein concentrations of the lysates were determined using BCA Protein Assay Kit (Pierce Biotechnology). Samples were heated at 95°C for 5 min in Laemmli sample buffer, and equal protein amounts were loaded on SDS-PAGE. Membranes were stained with primary antibodies for 16 hours at 4 °C. Then a secondary anti-rabbit or anti mouse-peroxidase antibodies (1:10,000; Jackson ImmunoResearch) were applied for 1 hour.

### Immunofluorescence

HeLa cells were washed with PBS, fixed with 4% paraformaldehyde (PFA), permeabilized with 0.5% Triton X-100 for 10 min and blocked with 10% FBS for 15 min. Fixed cells were stained with the indicated primary antibodies and subjected to fluorescently-tagged secondary antibodies staining (see antibody table). Finally, cells were mounted with Fluoromount-G (SouthernBiotech, Birmingham, AL) and imaged.

### Plasmids and antibodies

**Constructs:** GFP–α-tubulin and mCherry–α-tubulin were generated as previously described [11].

### Primary antibodies

Primary antibodies:
| Antigen | Source | Dilution | Company | Cat. Number | Used for |
| --- | --- | --- | --- | --- | --- |
| α-Tubulin | Mouse | 1:1000 | Sigma | T6199 | IF |
| β-Tubulin | Rabbit | 1:250 | abcam | ab6046 | IF |
| $\alpha$ and $\beta$ - Tubulin | Sheep | 1:500 | Cytoskeleton | #ATN02 | IF |
| IST-1 | Rabbit | 1:50 IF<br>1:1000 WB | Proteintech | 51002-1-AP | IF + WB |
| IST-1 | Mouse | 1:500 IF<br>1:5000 WB | Proteintech | 66989-1-Ig | IF + WB |
| CHMP2A | Rabbit | 1:50 IF<br>1:800 WB | Proteintech | 10477-1-AP | IF + WB |
| CHMP2B | Rabbit | 1:50 IF<br>1:800 WB | abcam | ab33174 | IF + WB |
| CHMP3 | Rabbit | 1:50 IF<br>1:500 WB | Sigma | SAB4301679 | IF + WB |
| CHMP1B | Rabbit | 1:750 IF<br>1:1500 WB | Sigma | HPA061997 | IF + WB |
| CHMP4B | Rabbit | 1:50 IF<br>1:800 WB | Proteintech | 13683-1-AP | IF + WB |

### Secondary antibodies

Anti-mouse Alexa 488/594/647 (1:1000, Life Technologies).

Anti-rabbit Alexa 488/594/647 (1:1000, Life Technologies).

Anti-sheep Alexa 594 (1:1000, Life Technologies).

### Live Cell Imaging

HeLa cells were plated at low density in four-well chamber slides (Ibidi, Martinsried, Germany), transfected 24 hours later with the indicated plasmids, and imaged at 24–48 hours post transfection. Z-stacks of selected cells with lower expression levels of the fluorescent proteins, undergoing cytokinesis were collected at the specified intervals, using a fully incubated confocal spinning-disk microscope (Marianas; Intelligent Imaging, Denver, CO), with a 63× oil objective (numerical aperture, 1.4), and were imaged on an sCMOS camera (Prime 95B, Teledyne Photometrics). Image processing and analysis were done using SlideBook version 6 (3I Inc).

### Structured Illumination Microscopy (SIM)

Cells were plated at low density on high-resolution #1.5 coverslips (Marienfeld, Lauda- Konigshfen, Germany) and fixed using 4% PFA. Cells were further subjected to immunostaining as described above. 3D SIM imaging of cells was performed using the ELYRA PS.1 microscope (Carl Zeiss MicroImaging). Thin z-sections (0.11–0.15 μm) of high-resolution images, were collected in three rotations and five phases for each channel.

Image reconstruction, channel alignment, and processing were performed in ZEN (Carl Zeiss MicroImaging), as described in [15, 25].

### Correlative Light-EM Microscopy (CLEM)

CLEM protocol for intercellular bridges was developed based on the protocol established by Avinoam et. al. for plasma membrane endocytosis [33].

#### Sample preparation

Cells were synchronized and grown on 3 mm Sapphire disks coated with carbon and then fixed with 3% PFA and 0.1% glutaraldehyde in a general cytoskeleton buffer (10 mM MES, 150 mM NaCl, 5 mM EGTA, 5 mM D-Glucose, 5 mM MgCl2, pH 6.1) for 15 min. After washing, samples were stained according to the standard Immunofluorescence protocol, and incubated in PBS supplemented with 10% BSA as a cryoprotectant. The samples were then mounted between carriers and rapidly vitrified by high-pressure freezing (Leica EM ICE). Freeze substitution was performed in 0.1% uranyl acetate in acetone at −90 °C using a Leica EM AFS2 system, followed by Lowicryl HM20 infiltration and UV polymerization at −25 °C.

#### Fluorescence Imaging of Resin Sections

Serial sections were cut at 350 nm thickness for transmission electron microscopy (TEM) and 1000 nm for scanning transmission electron microscopy (STEM) using a Leica Ultracut UCT ultramicrotome, mounted with a diamond knife (Diatome, Biel, Switzerland), and collected onto carbon-coated 200 mesh copper EM grids (Electron Microscopy Sciences, Hatfield, PA). After sectioning, grids were imaged using a spinning-disk confocal microscope (Marianas; Intelligent Imaging, Denver, CO) equipped with a 63× oil immersion objective (NA 1.4) and intracellular bridges were mapped.

#### Electron tomography

Mapped regions were subsequently imaged by electron microscopy (TEM or STEM) on a Thermo Fisher Scientific Talos F200C transmission electron microscope at 200 kV in TEM and STEM modes. The images in TEM mode were acquired using Ceta 16M CMOS camera and in STEM mode bright field (BF) detector was used for imaging. The tilt series were acquired with Thermo Fisher Scientific Tomography software (version 5.9) between angles varying from -60° to +60° with 1° step. Before each acquisition of an image with movement of the stage to the next tilt angle a cross correlation of tracking and focusing were made.

### Measurements

#### Bridge Diameter Calculations

HeLa cells transfected with GFP-Tubulin were imaged during cytokinesis, using a confocal spinning-disk microscope (Marianas; Intelligent Imaging) equipped with a 63× oil objective. The diameter of the intercellular bridge was measured from the acquired time-lapse images, using SlideBook version 6 (3I Inc). Measurements were taken from the rims of the DZ, using a line intensity profile perpendicular to the bridge axis, as described in Gershony et. al. [34]

#### Image segmentation

HeLa WT and CHMP2A KO cells were fixed, permeabilized, and stained with WGA (membrane), CHMP3 (cytosolic), and Hoechst (nuclei). Samples were imaged using a spinning disk confocal microscope (as described above; Fig. S2A). Cell and nuclei segmentations were performed using a python script using OpenCV and SK-Image packages based on Hoechst and WGA staining (nuclei and cells, respectively) [35]. Area of segmented regions were then measured and plotted (Fig. S2).

### Statistical analysis

Statistical analysis was performed using Graph Pad Prism version 9.00 for Windows (La Jolla, CA, USA). For comparisons between two groups, a two-tailed unpaired t-test was used when the data followed a normal distribution. If the data did not follow a normal distribution, a Mann-Whitney U test was applied. Data was considered statistically significant at *p* < 0.05. Statistical significance for the distribution of localization phenotypes (Fig. 3C) was calculated using Fisher’s exact test.

## ACKNOWLEDGMENTS

We thank all members of the Elia lab for critical feedback throughout the project and for reading the manuscript. This research is funded by the Israeli Science Foundation (ISF) NE Grant no. 1323/18.

## AUTHOR CONTRIBUTIONS

NK and NE conceptualized the project. NK performed and analyzed all experiments. NE wrote the manuscript with the help of NK. DN provided technical help. AU provided technical help with EM data acquisition. SK and OA assisted in developing CLEM protocol. IH performed image segmentation analysis. All authors read and revised the manuscript.

## Competing Interest Statement

We declare no competing interests.

---

**Fig. S1.**
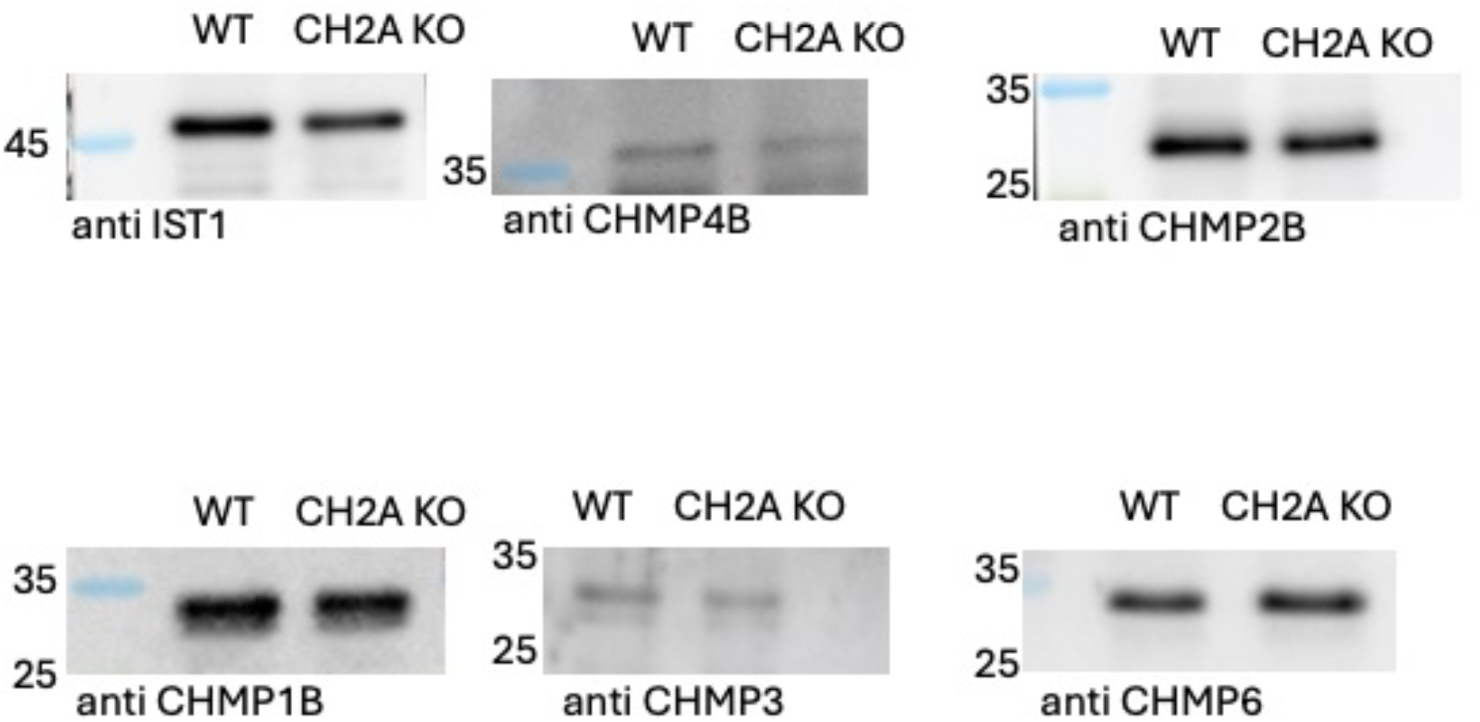
Comparative analysis of protein expression in WT and CHMP2A KO HeLa cells. Representative Western blotting analysis showing the levels of IST1, CHMP4B, CHMP2B, CHMP1B, CHMP3 in whole cell lysates from wild-type (WT) and CHMP2A KO cells. Equal amounts of total protein (30 μg per lane) were loaded for each sample. Protein bands indicate similar expression levels between the two conditions.

**Figure S2.**
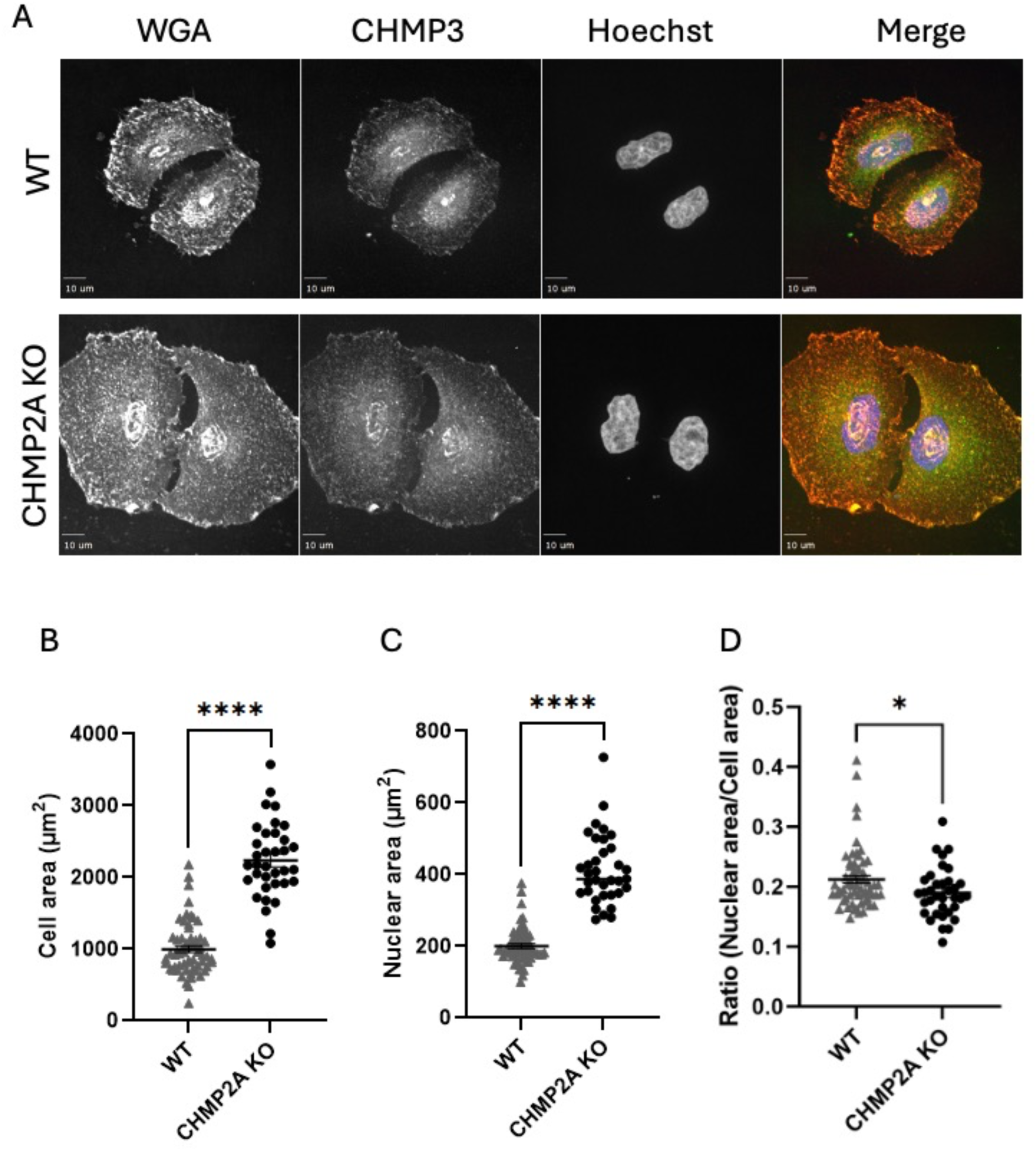
Quantitative analysis of cell size in WT and CHMP2A KO cells. (**A-D**) HeLa WT (n=60) and CHMP2A KO (n=35) cells were stained with WGA (membrane), CHMP3 (ESCRT-III) and Hoechst (nuclei) and imaged using spinning-disk confocal microscopy. Cell and nuclear were segmented based on WGA and Hoechst staining respectively and areas of segmented regions were calculated (see methods). Note, that both cell area and nuclear area measurements show a significant enlargement in CHMP2A KO cells compared to WT cells. The nuclear-to-cell area ratio remains comparable between cell types, indicating proportional scaling of nuclear and cytoplasmic size in CHMP2A KO cells. Each dot represents an individual cell. Data are presented as mean ± SD. Statistical significance was assessed using t-test *p*<0.05(\**), p < 0.001 (\*\*\*\**).

**Fig. S3.**
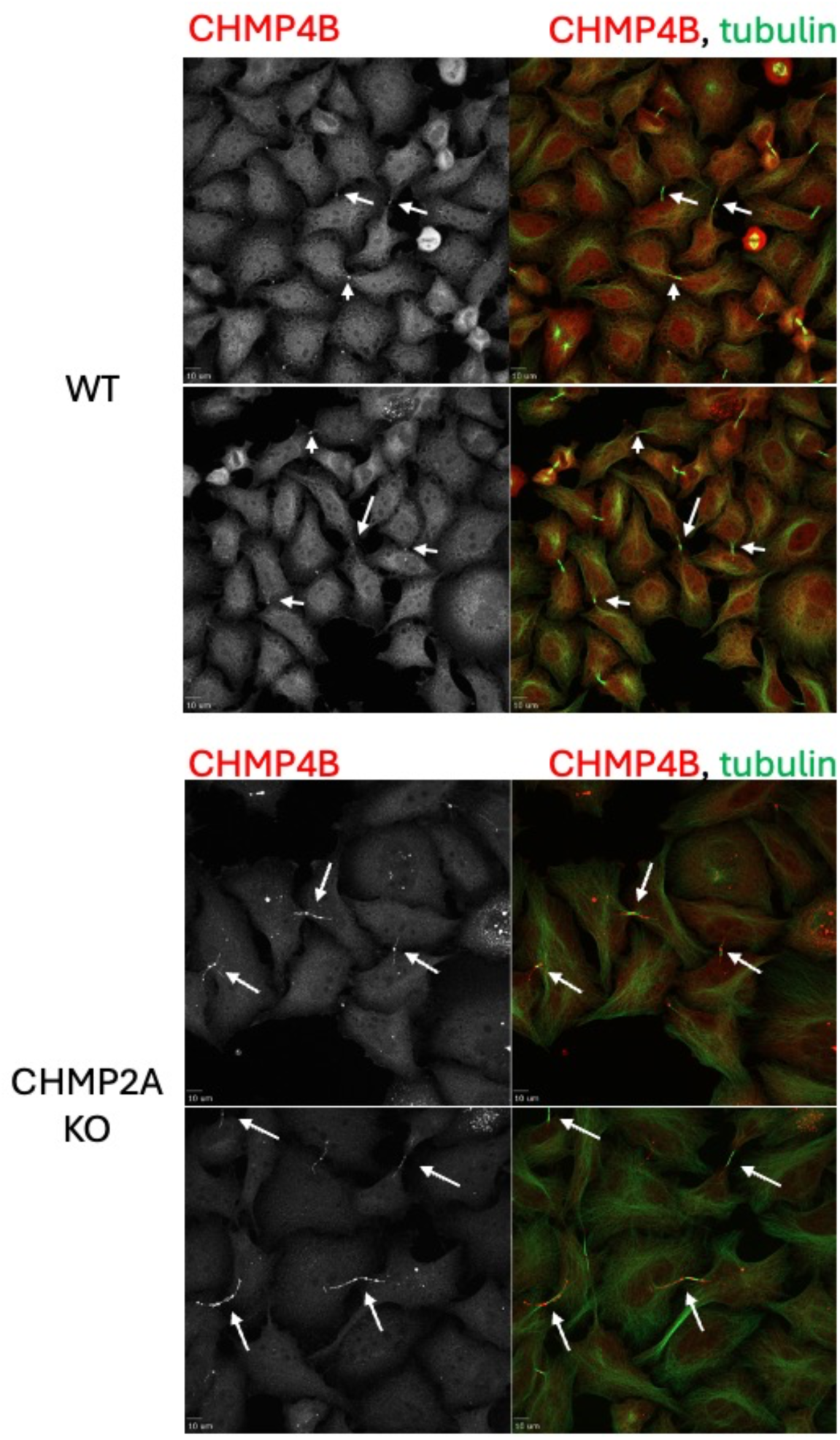
The No DZ localization is a frequent phenotype of CHMP4B localization in CHMP2A KO cells. Representative maximum projections confocal images of synchronized HeLa cells (WT, top; CHMP2A, bottom), stained with α–tubulin (green) and α-CHMP4B (red) antibodies. Arrows indicate intercellular bridges. Note the abundance of the ‘No DZ localization’ phenotype of CHMP4B in CHMP2A KO cells, manifested by a string-like pattern along the intercellular bridge.

**Figure S4.**
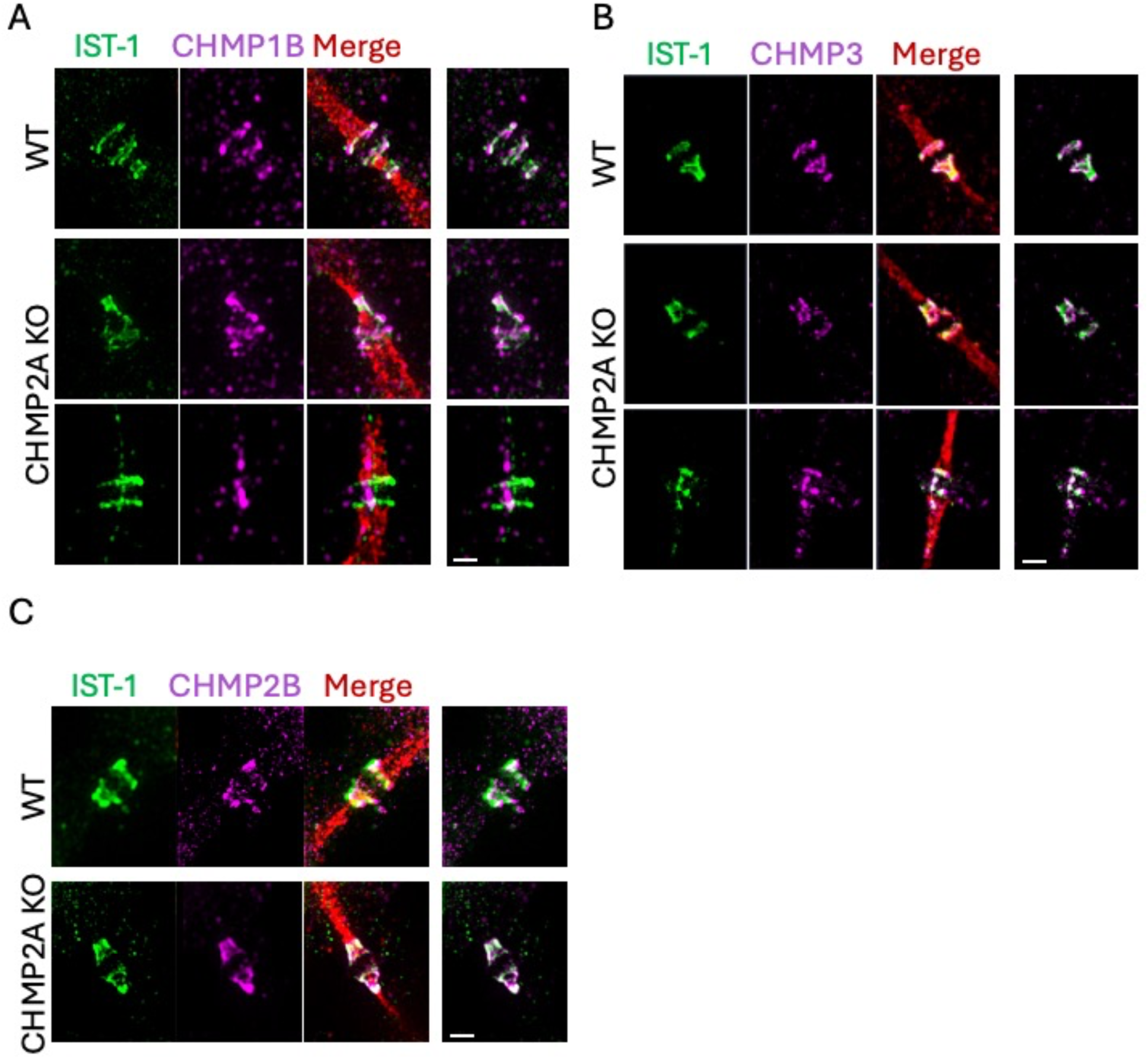
Differential localization patterns of ESCRT-III protein pairs upon CHMP2A depletion within individual cells. (A-C) Representative SIM images of synchronized HeLa cells (WT and CHMP2A KO). HeLa WT cells (top panels) or CHMP2A KO cells (bottom panels) were labeled with α-- IST1 (green, left), and α--ESCRT-III (as indicated, magenta, second) and α-tubulin (red). The overlayed image is presented on the third panel. Right panel, overlaid images of ESCRT-III proteins highlighting the coordinated / uncoupled assembly of the labeled proteins. Data corresponds to matrixes shown in Fig. 5B. Scale = 1 μm.

## Notes

### Competing Interest Statement

The authors have declared no competing interest.

